# The two sides of resistance: aggressiveness and mitotic instability as the Achilles’ heel of Osimertinib-resistant NSCLC

**DOI:** 10.1101/2025.09.25.678204

**Authors:** Paolo Armanetti, Eva Cabrera San Millan, Raffaella Mercatelli, Chiara Cavallini, Alice Chiodi, Ettore Mosca, Arianna Consiglio, Nicola Losito, Luca Menichetti, Emilia Bramanti, Azhar Ali, Elena Levantini, Giorgia Maroni

## Abstract

Non-small cell lung cancer (NSCLC) represents majority of lung cancer cases and remains a leading cause of cancer mortality worldwide. Tumors carrying activating mutations in the epidermal growth factor receptor (EGFR) are highly sensitive to EGFR tyrosine kinase inhibitors (TKIs), with third-generation inhibitors such as Osimertinib now established as standard of care. However, acquired resistance to Osimertinib inevitably develops, involving both genetic and non-genetic mechanisms, the latter playing a major role in sustaining cellular plasticity and promoting tumor aggressiveness. Among regulators of adaptive programs, the Polycomb protein BMI1 has emerged as a key factor driving stemness, epithelial-to-mesenchymal transition (EMT), and therapy resistance in multiple cancers, yet its role in Osimertinib resistance remains poorly defined.

Here, we show that Osimertinib-resistant H1975 cells, which display greater aggressiveness than their parental counterparts, are enriched in BMI1 target genes and mitotic cell-cycle pathways, establishing a dependency on microtubule dynamics and mitotic control. Functionally, BMI1 drives migration, invasiveness, and tumor progression in resistant cells. This mitotic dependency creates a therapeutic vulnerability that can be exploited with Unesbulin (PTC596), a BMI1 inhibitor that destabilizes microtubules and induces mitotic catastrophe, thereby effectively suppressing tumor growth *in vitro* and *in vivo*.

Our findings establish BMI1 as a central mediator of Osimertinib resistance and provide a mechanistic and therapeutic rationale for targeting BMI1 and mitotic weaknesses in refractory EGFR-mutant NSCLC.

## Introduction

Non-small cell lung cancer (NSCLC) accounts for ∼85% of lung cancer cases worldwide and remains a leading cause of cancer-related mortality (O’Leary et al., 2020). Tumors harboring activating mutations in the epidermal growth factor receptor (EGFR) represent a clinically relevant subgroup, characterized by high sensitivity to EGFR tyrosine kinase inhibitors (TKIs) (Levantini et al., 2022). Third-generation TKIs such as Osimertinib, initially developed to overcome T790M-mediated resistance, are now the standard of care for first-line settings due to their ability to target both sensitizing and resistance EGFR mutations (Levantini et al., 2022).

Despite initial responses, the development of acquired resistance to Osimertinib is inevitable, arising typically within 1-2 years of treatment initiation (Oxnard et al., 2018). Several mechanisms have been described, including secondary EGFR mutations (e.g., C797S), activation of bypass signaling pathways, histological transformation, and engagement of epithelial-to-mesenchymal transition (EMT) programs (Oxnard et al., 2018). We already reported a mitochondria-centered adaptive program where mitochondrial remodeling and activation of the pyruvate-acetaldehyde-acetate (PAA) pathway converge to promote cell survival and drug tolerance in Osimertinib resistant NSCLC (Maroni et al., 2025 *BioRxiv* https://doi.org/10.1101/2025.06.27.661913, Maroni et al., 2025 *BioRxiv* https://doi.org/10.1101/2025.07.04.663159). Non-genetic adaptive mechanisms are increasingly recognized as key contributors to resistance, supporting cellular plasticity and enabling tumor survival under therapeutic pressure. However, the functional consequences of such adaptations, particularly in relation to tumor aggressiveness and invasive potential, remain incompletely understood.

Polycomb group proteins, particularly BMI1, a core component of the Polycomb Repressive Complex 1 (PRC1), regulate stemness, EMT, DNA damage response, and therapy tolerance in multiple cancers (Fitieh et al., 2021; Ren et al., 2016; Yang et al., 2019). In NSCLC, elevated BMI1 expression has been associated with poor prognosis, enhanced metastatic potential, and resistance to chemo- and radio-therapy (Levantini et al., 2022; Maroni et al., 2021; Maroni et al., 2024; Meng et al., 2012; Yong et al., 2016; Zhang et al., 2017). Its role in Osimertinib-resistant EGFR-mutant NSCLC, however, remains poorly understood.

Here, we investigated Osimertinib-resistant H1975 cells (H1975 Osi-R) and observed an enrichment of BMI1 target genes and mitotic pathways, including spindle assembly, G2/M checkpoint, Myc and E2F transcriptional programs, and DNA repair processes. This transcriptional landscape suggests that resistant cells acquire dependency on cell-cycle and mitotic regulation, as well as microtubule dynamics, potentially predisposing them to mitotic stress and genomic instability. Such vulnerabilities could be therapeutically exploitable, particularly in the context of emerging strategies targeting BMI1 and mitosis-associated processes, mediated by microtubules.

Unesbulin (PTC596), a small-molecule BMI1 inhibitor, has been evaluated in a phase I clinical trial and was well tolerated in patients with advanced solid tumors (PMID: 33440067). Preclinical studies indicate that Unesbulin destabilizes microtubules, impairs mitotic progression, and triggers apoptotic cell death (Jernigan et al., 2021; Nishida et al., 2017; Shapiro et al., 2021). A phase II/III trial has been registered to assess its safety and efficacy (EudraCT 2022-000073-12). Given the reliance of Osimertinib-resistant cells on BMI1 signaling and mitotic pathways, Unesbulin represents a promising candidate to exploit these vulnerabilities.

Here, we investigated the biological features of H1975 Osi-R cells and tested the efficacy of Unesbulin both *in vitro* and *in vivo*. We demonstrate that BMI1 acts as a key mediator of Osimertinib resistance, linking/associate with it to increased migratory behavior and aggressive transcriptional programs, and that pharmacological inhibition of BMI1 with Unesbulin effectively suppresses tumor progression. These findings provide a strong rationale for therapeutic targeting of BMI1 and mitotic vulnerabilities in EGFR-mutant NSCLC that has acquired resistance to Osimertinib.

## Results

### H1975 Osi-R cells exhibit higher migration and mitotic dependency

To investigate the mechanisms underlying Osimertinib resistance, we used previously generated H1975 cells resistant to Osimertinib (Osi-R) (Maroni et al., 2025 *BioRxiv* https://doi.org/10.1101/2025.06.27.661913), and compared their *in vitro* behavior with parental counterparts (Par). Since H1975 OR cells exhibit a metabolic shift (Maroni et al., 2025 *BioRxiv* https://doi.org/10.1101/2025.06.27.661913, Maroni et al., 2025 *BioRxiv* https://doi.org/10.1101/2025.07.04.663159), we first assessed whether this altered metabolism impacted proliferation. BrdU incorporation assays showed no significant differences in proliferation between Osi-R and parental cells at 24 hours (Fig. 1A), indicating that resistance does not involve increased proliferative ability.

**Figure 1.**
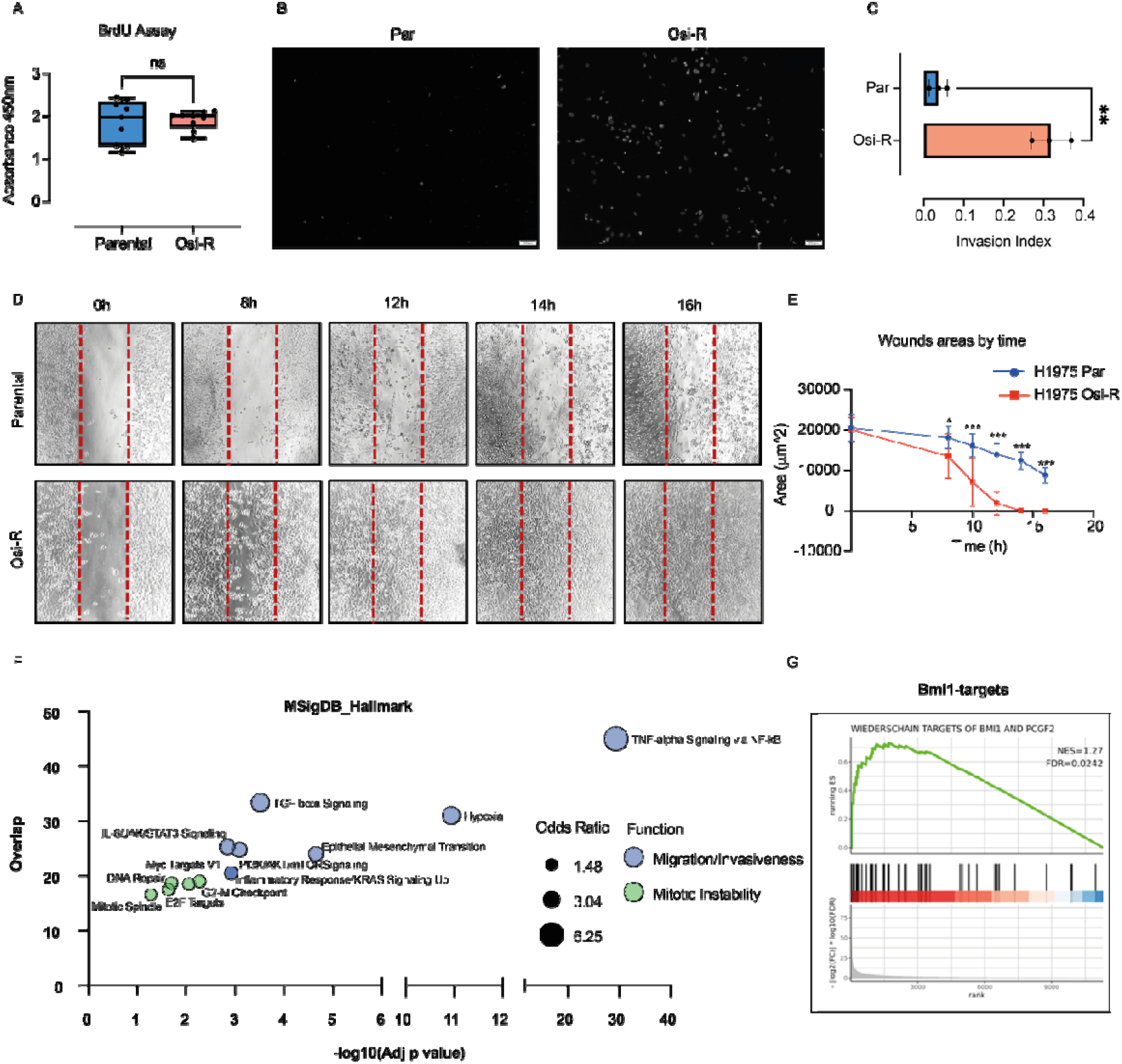
Migration assays and transcriptomic analysis reveal enhanced invasiveness and mitotic dependency in Osimertinib-resistant EGFR mutant NSCLC cells. (a) BrdU cell proliferation assays in H1975 Par (blue) and Osi-R (red) cells. Error bars represent standard deviation (SD); no significant differences in proliferation were detected (ns, not significant, Welch’s T test, n=9). (b) Transwell migration assay images of H1975 cells. DAPI staining of the nuclei in Par cells (left panel) and in Osi-R H1975 cells (right panel). (c) Quantitative analysis of the transwell migration assay of H1975 Par (blue) and Osi-R (red) cells. Error bars represent standard deviation (SD); p-value is indicated (**p<0.0021, Welch’s T test). (d) Wound-healing assay images of Par (top panel) and Osi-R (bottom panel) H1975 cells at the indicated time points. (e) Quantitative analysis of wound healing assay of H1975 Par (blue) and Osi-R (red) cells. Error bars represent standard deviation (SD); p-value are indicated (*p<0.032, **p<0.0021, ***p<0.0002, Multiple Mann-Whitney Tests). (f) RNA-seq pathway enrichment analysis (EnrichR, MSigDB_Hallmark) showing enrichment statistically significant of pathways associated with migration /invasiveness (blue) and mitotic instability(green) in Osi-R cells. Dot size indicates odd ratio. (g) Gene set enrichment analysis (GSEA) of BMI1 target genes showing positive enrichment in Osi-R cells Normalized Enrichment Score (NES) = 1.27; FDR: 0.0242).

Despite unchanged proliferation, Osi-R cells displayed a more aggressive phenotype. Transwell migration assays showed enhanced migratory capacity (Fig.1B), as indicated by DAPI staining, which showed a higher number of migrated Osi-R cells compared to parental controls (Fig. 1B). Quantification confirmed a ∼8.8-fold increase (Osi-R=0.3179, Par=0.036) in the migration index (Fig. 1C). Wound healing assays further confirmed their heightened invasiveness in that H1975 Osi-R cells started repopulating the gap as early as 8 hours after scratching (Fig. 1D-E), and displayed complete closure within 14 hours (Fig. 1D-E).

RNA-seq pathway analysis (EnrichR, MSigDB_Hallmark database), using our data previously reported (Maroni et al., 2025 *BioRxiv* https://doi.org/10.1101/2025.06.27.661913), demonstrated enrichment of pathways associated with invasiveness, including TNF-alpha Signaling via NF-kB, Hypoxia, TGF-beta Signaling and Epithelial Mesenchymal Transition, PI3K/AKT/mTOR Signaling, Inflammatory Response, KRAS Signaling Up and IL-6/JAK/STAT3 Signaling, in Osi-R cells (Fig 1F).

In parallel, pathways related to mitotic regulation (mitotic spindle, G2M checkpoint, E2F targets, Myc targets, DNA repair were also enriched in OR cells (Fig. 1F), indicating both enhanced mitotic activity, potential mitotic instability and dependency on microtubule dynamics.

GSEA revealed a positive enrichment in Osi-R cells for BMI1 target genes (Normalized Enrichment Score (NES) = 1.27; FDR: 0.0242) (Fig. 1G), linking resistance to BMI1-driven transcriptional changes.

### Unesbulin induces apoptosis of H1975 Osi-R cells *in vitro*

Given the reprogramming of Osi-R cells, which includes dependency on BMI1 and mitotic processes, we investigated their response to Unesbulin (1 μM), a tubulin-binding agent with BMI1 inhibitory activity.

Remarkably, cytotoxic response was present at already 24 hours post-treatment, with majority of cells displaying extensive cell death (Fig. 2A).

**Figure 2.**
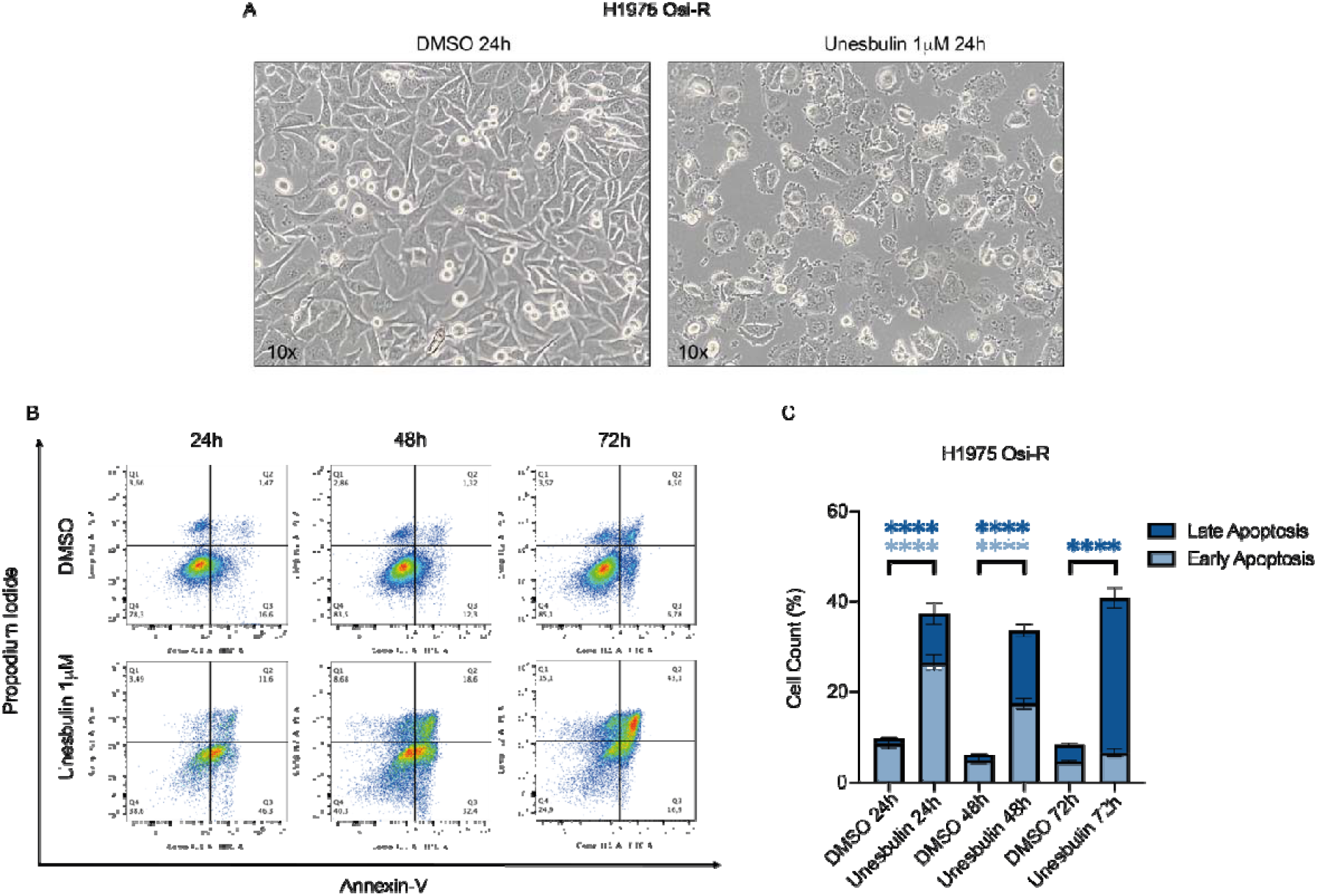
Unesbulin treatment induces apoptosis in Osimertinib-resistant EGFR mutant NSCLC cells. (a) Representative image of H1975 Osi-R cells treated with DMSO (control, left panel) or Unesbulin (right panel) after 24 h. Images were acquired at 10× magnification. (b) Annexin V/PI flow cytometry analysis of H1975 Osi-R cells treated with DMSO or Unesbulin for 24, 48, and 72 hours. Plots show progressive increases in Annexin V+ (FITC-A channel) (early apoptotic) and Annexin V+/PI+ (PE-A channel) (late apoptotic) populations in Unesbulin-treated cells. (c) Quantitative analysis of apoptotic populations from flow cytometry data in Unesbulin-treated versus DMSO-treated cells. Early apoptosis is shown in light blue, and late apoptosis in dark blue. Error bars represent standard deviation (SD); p-values are indicated (****p<0.0001, 2-way ANOVA)

Annexin-V/Propidium Iodide (PI) FACS-based analysis confirmed apoptosis. Specifically, after 24 hours of Unesbulin treatment, Osi-R cells showed a significant ∼3-fold increase in Annexin-V^+^ cells as compared to DMSO-treated controls (p: 8.3×10^-14^) (Fig. 2B-C), indicating early apoptotic induction. At 48h and 72h, a progressive rise in double positive (Annexin-V^+^/PI^+^) cells was observed (16% and 34%, respectively, Fig. 2B-C), consistent with increased late-stage apoptosis relative to controls (p: 4×10^12^ and 2×10^14^, respectively, Fig. 2B-C). these data demonstrate that Osi-R cells are sensitive to Unesbulin treatment *in vitro*.

### Unesbulin causes tumor regression of H1975 Osi-R xenografts *in vivo*

Next, we evaluated the *in vivo* growth capacity of H1975 resistant cells compared to their parental counterparts. Both cell types were transplanted into immunocompromised mice, and tumor growth (mm^3^) was monitored weekly by ultrasound (US) starting on day 10 post-implantation. Over a 3-week period, ultrasound measurements revealed no significant difference in growth rates between Osi-R and Par-derived xenografts (Fig. 3A), consistent with the BrdU proliferation assays in culture.

**Figure 3.**
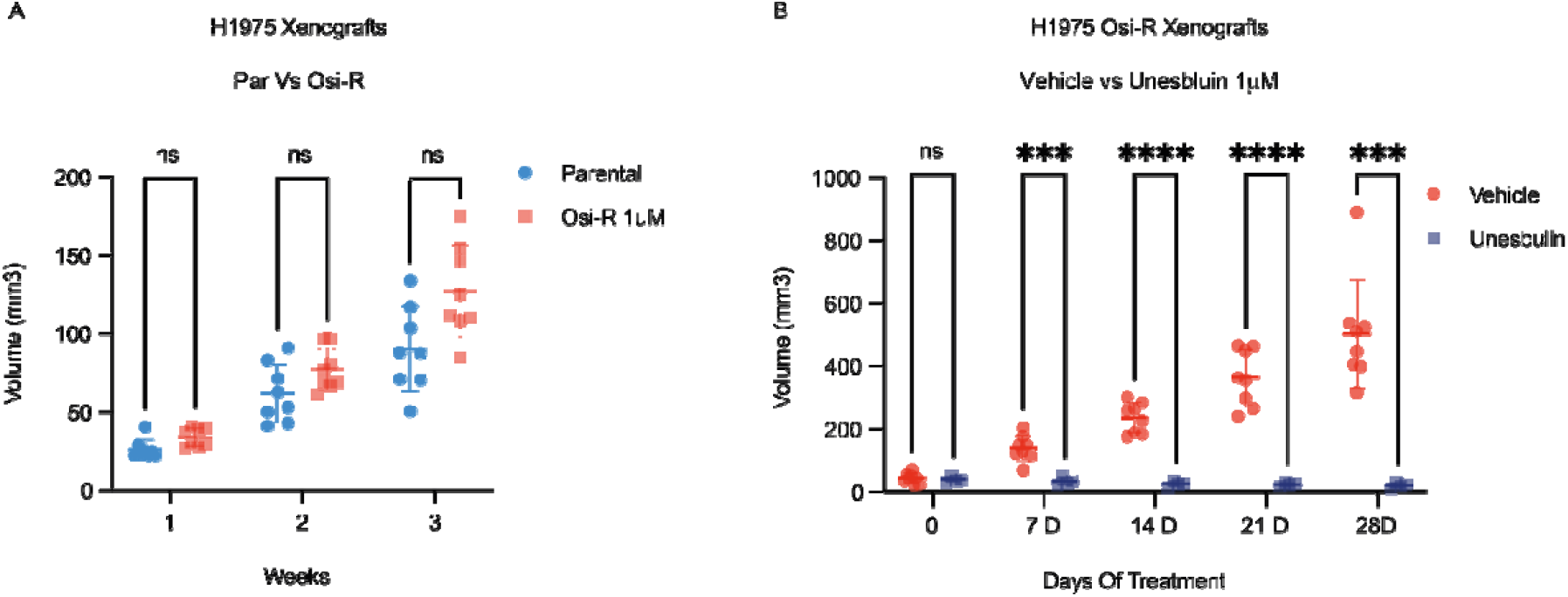
Unesbulin treatment induces tumor regression in Osimertinib-resistant EGFR mutant NSCLC cells *in vivo*. (a) Tumor growth kinetics of H1975 Par (blue, n=8) and Osi-R (red, n=8) xenografts. Error bars represent standard deviation (SD); no significant differences were detected (ns, not significant). (b) Tumor growth kinetics of H1975 Osi-R treated with Vehicle (red, n=8) or Unesbulin (blue, n=4). Error bars represent standard deviation (SD); p-values are indicated (ns, not significant, ***p<0.0002, ****p<0.0001, Multiple Unpaired T tests).

Once Osi-R-derived subcutaneous tumors reached ∼40 mm^3^, we initiated biweekly Unesbulin treatment (12mg/Kg). Strikingly, while vehicle-treated tumors grew to an average final volume of ∼500 mm^3^, Unesbulin-treated tumors shrank to ∼20 mm^3^. Such 25-fold difference in volume (p=3.5×10^-5^), demonstrates that Unesbulin effectively targets the vulnerabilities of OR tumors *in vivo* (Fig. 3B).

### Bmi1 upregulation in H1975 parental cells resembles the Osi-R phenotype both *in vitro* and *in vivo*

To functionally assay BMI1 contribution to driving resistance-associated features, we overexpressed it in H1975 parental cells. Western blot analysis confirmed BMI1 overexpression in cells transfected with a pCMV3-BMI1 construct (Bmi1-up), compared to those transfected with the empty pCMV3 control vector (Fig. 4A). BrdU assays after 24 h revealed no significant difference in proliferation between Bmi1-up and pCMV3 control cells (Fig. 4B).

**Figure 4.**
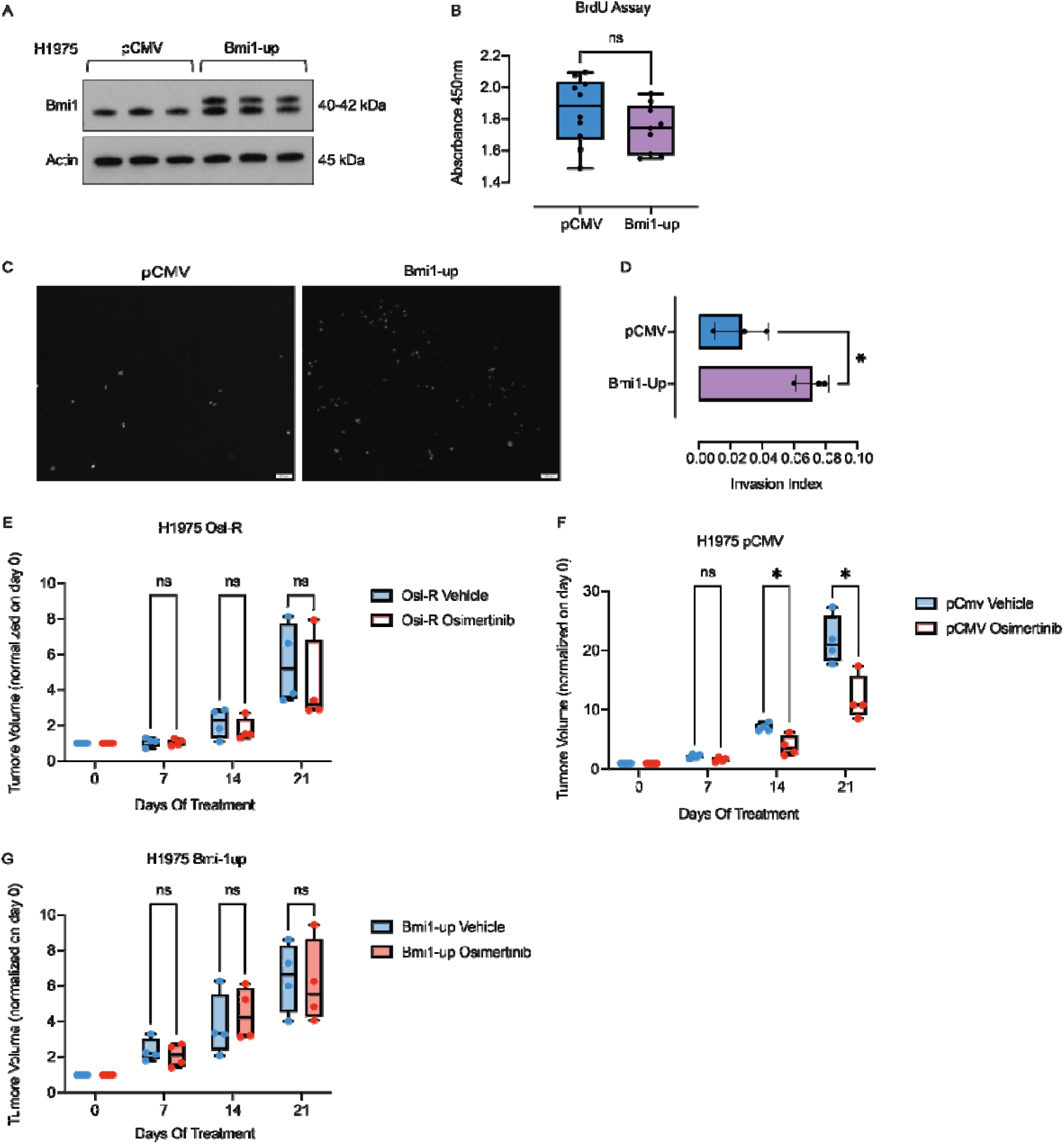
BMI1 overexpression enhances migration and confers resistance to Osimertinib in EGFR mutant NSCLC cells. (a) Western blot analyses of H1975 pCMV3 and pCMV3-BMI1up. Protein lysates were immunoblotted with an antibody against BMI1. β-actin was used as a loading control. Expected molecular weights (kDa) are indicated. (b) BrdU cell proliferation assays in H1975 pCMV3 (blue) and pCMV3-BMI1up (purple). Error bars represent standard deviation (SD); no significant differences in proliferation were detected (ns, not significant, Welch’s T test, n=10). (c) Transwell migration assay images of H1975 cells. DAPI staining of the nuclei in H1975 pCMV3 cells (left panel) and in H1975 pCMV3-BMI1up cells (right panel). (d) Quantitative analysis of the transwell migration assay of H1975 pCMV (blue) and pCMV3-BMI1up (purple) cells. Error bars represent standard deviation (SD); p-value is indicated (*p<0.0332, Welch’s T test). (e–g) Tumor growth kinetics of xenografts with H1975 Osi-R (e), H1975 pCMV3 (f), and H1975 pCMV3-BMI1up (g) cells treated with vehicle (blue) or Osimertinib (red). Tumor volume ratios were normalized to day 0. Error bars represent standard deviation (SD); p-values are indicated (ns, not significant; *p<0.0032, Multiple Unpaired T Tests, n=4).

Notably, transwell migration assays showed a ∼3-fold increase in migrating Bmi1-up cells compared to pCMV3 controls (p=0,0246, Fig. 4C-D), indicating an acquired enhancement in migratory ability upon BMI1 overexpression in P cells.

Next, we tested the impact of BMI1 overexpression in *in vivo* settings. To this aim, we compared tumor growth response to Osimertinib treatment in xenografts derived from H1975 Osi-R, Bmi1-up, and pCMV3 cells. Mice received biweekly drug treatment, and tumor growth was monitored for up to 3 weeks.

As proof-of-concept, H1975 Osi-R xenografts were resistant to Osimertinib (10mg/Kg), showing no reduction in tumor growth compared to vehicle-treated Osi-R tumors (Fig. 4E). As expected, xenografts derived from pCMV3-transfected cells (reflecting Osimertinib-sensitive parental H1975 cells), responded to Osimertinib, showing a significant reduction in tumor volume as early as 2 weeks post drug treatment (Fig. 4F). Notably, Bmi1-up xenografts displayed resistance to Osimertinib, growing comparably to vehicle-treated tumors (Fig. 4G).

Together, our data demonstrate that BMI1 overexpression is *per se* sufficient to confer *in vivo* drug resistance, while simultaneously enhancing migratory behavior, positioning BMI1 as a central node linking resistance and aggressiveness.

## Discussion

Our study defines the biological and molecular features/landscape of NSCLC cells that acquire resistance to Osimertinib, with a focus on the contribution of BMI1 signaling to the resistant phenotype. We show that H1975 OR cells do not display altered proliferative rates in culture or *in vivo* when compared with parental counterparts. Instead, they undergo a profound phenotypic shift characterized by enhanced migratory and invasive potential, coupled with a strong reliance on mitotic machinery, hallmarks of a more aggressive behavior.

Even if proliferation rates remain unchanged, the migratory phenotype and molecular rewiring of H1975 OR cells suggest a selective advantage under therapeutic pressure, enabling tumor progression despite drug treatment. This observation reinforces the emerging concept that resistance is not always driven by increased proliferative fitness but also by phenotypic plasticity and adaptive signaling programs that sustain survival and dissemination (Gupta et al., 2019).

Consistently, transwell and wound-healing assays demonstrated markedly increased migratory ability of resistant cells. This functional observation was supported by RNA-seq-based pathway analyses that revealed enrichment of networks known to promote invasion and EMT, including TGF-β, NF-κB, IL6/JAK/STAT3, and PI3K/AKT/mTOR signaling (Kartikasari et al., 2021; Wong et al., 2022). The co-enrichment of hypoxia- and inflammation-related pathways further suggest that transcriptional rewiring fosters tumor plasticity and adaptive responses. This is in line with recent evidence indicating that resistance can arise through non-genetic mechanisms, particularly phenotypic switching (Chen et al., 2023). For instance, EMT-like reprogramming has been linked to drug tolerance, collective migration in metastasis, and immune evasion, thereby providing a strong survival advantage under hostile microenvironmental conditions. In parallel, the enrichment of mitotic spindle, G2M checkpoint, Myc and E2F targets, as well as DNA repair signatures identified in OR cells suggest that resistant cells acquire a dependency on cell-cycle regulation and microtubule dynamics to cope with mitotic stress and to maintain genomic stability. Cancer cells’ dependence on mitotic machinery, a hallmark of therapy-induced adaptation (Sarkar et al., 2021), provides transient survival advantages yet simultaneously creates a pharmacologically actionable *Achilles’ heel*.

Pathway and GSEA analysis also identify significant enrichment in BMI1 target genes, further validating its role in transcriptional reprogramming and therapy escape. Notably, BMI1 overexpression alone in parental cells is sufficient to reproduce/recapitulate both increased migration *in vitro* as well as Osimertinib resistance *in vivo*, pointing to BMI1 as a central node linking resistance acquisition with enhanced invasiveness. This aligns with prior evidence implicating BMI1 in stemness, EMT induction, and drug tolerance in disparate tumor contexts (Fitieh et al., 2021; Ren et al., 2016; Yang et al., 2019). Our data extend this role to EGFR-mutant NSCLC, positioning BMI1 as both a biomarker and a therapeutic vulnerability in Osimertinib resistance. Importantly, resistance generates collateral dependencies on mitotic spindle dynamics and G2/M checkpoint control, rendering OR cells sensitive to pharmacological perturbation.

Importantly, Unesbulin capitalizes on these vulnerabilities through a dual mechanism of action: Unesbulin lowers BMI1 protein levels, disrupting BMI1-driven transcriptional programs and disrupt tubulin polymerization, destabilizing microtubules and leading to spindle dysfunction. This combination triggers rapid and extensive apoptotic cell death *in vitro* and drives regression of OsiR xenografts *in vivo*, demonstrating a therapeutic strategy that attack the weaknesses created by resistance itself.

The lethality of Unesbulin in this context likely reflects the fact that Osimertinib-resistant cells are already “stressed” by their dependence on mitotic spindle dynamics and G2/M checkpoint control. Additional perturbation of microtubule integrity by Unesbulin shifts the balance toward catastrophic mitotic failure. Thus, Unesbulin does not merely suppress a key resistance driver (BMI1) but also exploits the collateral vulnerabilities that resistance itself creates.

Together, these findings identify BMI1 as a mediator of resistance and migration and establish Unesbulin as a promising therapeutic option for EGFR-mutant NSCLC that has acquired resistance to Osimertinib. Clinically, these results support further evaluation of BMI1 inhibition both as a salvage strategy in resistant tumors and as a rational combination with Osimertinib to delay or prevent the emergence of BMI1-driven, mitotically dependent subclones. Such strategy may ultimately extend the durability of TKI responses and improve patient outcomes.

## MATERIAL AND METHODS

### Cell culture conditions

The parental human lung cancer cell line H1975, which harbors the EGFR L858R/T790M mutation, was obtained from ATCC. The osimertinib-resistant variant was generated by gradually exposing H1975 parental cells to increasing concentrations of osimertinib, up to a final concentration of 1 μM, at which the cells remained viable. Cells were maintained in RPMI-1640 medium supplemented with 10% fetal bovine serum (FBS), 1% L-glutamine, and 1% penicillin/streptomycin, at 37°C in a humidified incubator with 5% CO2.

### Generation of H1975 pCMV3-BMI1up and pCMV3 cells

H1975 cells were transfected with pCMV3 backbone or containing BMI1 cDNA using Fugene HD (Promega) according to manufacturer’s protocol. 48 hours after transfection, cells were into medium containing hygromycin B at 200µg/ml with fresh medium replacement every 4 days. After 2 weeks, surviving cells were observed and single cells resistant to hygromycin B were transferred into 96-well plates to ensure that they could grow into hygromycin B resistant colonies. BMI1 expression was validated by determining protein and mRNA expressions.

### Enrichment Analysis

The list of significantly upregulated genes was analyzed using the EnrichR platform for functional enrichment (Chen et al., 2013; Kuleshov et al., 2016; Xie et al., 2021). Enrichment analysis was performed using MSigDB_Hallmark database, to identify overrepresented biological pathways and cellular functions. GSEA was performed on the list of genes ranked by the product between absolute log2 fold change (Osi-R vs Par) and adjusted p-value (Ritchie et al., 2015), corrected through the Benjamini-Hocberg procedure (Benjamini et al., 2001), using R software environment.

### Cell Collection and Protein Extraction

H1975 pCMV3-BMI1up and pCMV3 cells were detached from culture plates using trypsin-EDTA, centrifuged at 261x g for 5 minutes at room temperature, and washed with PBS. The resulting cell pellet was snap-frozen in liquid nitrogen and stored at −80°C until further processing. To extract proteins, the cell pellet was lysed for 30 minutes in 1X Triton X-100 (Thermo Scientific, #A16046) supplemented with cOmplete EDTA-free Protease Inhibitor Cocktail and PhosStop Phosphatase Inhibitor Cocktail. The lysates were incubated on ice for 30 minutes with vortexing every 10 minutes. After incubation, the samples were centrifuged at 12,000 × g for 20 minutes at 4°C, and the supernatant was collected and stored at −80°C.

### SDS-PAGE and Western Blotting

For protein analysis, 15 μg of protein from each sample (H1975 pCMV3-BMI1up and pCMV3 cells) was separated by SDS-PAGE on 10% gel and subsequently transferred to nitrocellulose membranes with the TransBlot Transfer System (Bio-Rad). Membranes were blocked for 1 hour at room temperature in either 5% non-fat dry milk or 5% bovine serum albumin (BSA) diluted in Tris-buffered saline with 0.1% Tween-20 (1X TBS-T). Primary antibodies (listed below) were incubated overnight at 4°C. After incubation, membranes were washed three times for 10 minutes each with 1X TBS-T at room temperature. HRP-conjugated secondary antibodies (listed below) were then applied for 1 hour at room temperature. Proteins bands were detected by chemiluminescence (Pierce ECL Western Blotting Substrate Cat #32106) and visualized using the Chemidoc Imaging System (Biorad). Membranes were subsequently stripped using Stripping Buffer Solution (Himedia, #ML163), as per the manufacturer’s instructions, and re-probed with an anti-β-actin mouse antibody to verify equal protein loading. After three washes with 1X TBS-T, membranes were incubated with the appropriate anti-mouse secondary antibody (details below).

| **Antibodies and dilution** |  |
| --- | --- |
| **Primary antibody** | **Dilution** |
| anti-β-actin mouse antibody (Santa Cruz # C3022) | 1/1000 (in 5% non fat milk) |
| anti-BMI1 rabbit antibody (Cell Signaling Technologies #6964) | 1 /1000 (in 5% non fat milk) |
| **Secondary antibody** | **Dilution** |
| anti mouse (Immunoreagents # NC27606) | 1 2000 (in 5% non fat milk) |
| anti rabbit (Cell Signaling Technologies #7074) | 1 2000 (in 5% non fat milk or BSA) |

### BrDU cell proliferation assay

Cell proliferation was assessed using the BrdU Cell Proliferation Assay Kit (Sigma Aldrich, Cat. #QIA58) according to the manufacturer’s instructions. Briefly, H1075-Par and H1075-OsiR, pCMV3-BMI1up and pCMV3 cells were seeded in 96-well plates and allowed to adhere for 24 h until they reached approximately 40–60% confluence. BrdU was added and incubated for 2–24 h at 37°C in a humidified incubator with 5% CO_2_, after which cells were fixed. Incorporated BrdU was detected with mouse anti-BrdU antibody (1:1000), followed by HRP-conjugated anti-mouse IgG secondary antibody (1:1000). Substrate solution was added and incubated for 15 minutes at room temperature and then reaction was stopped with the stop solution. Absorbance was measured at 450 nm using a spectrophotometric plate reader.

### Transwell migration assays

Transwell migration assays was performed using Boyden Chamber from Corning® BioCoat™ Matrigel® Invasion Chambers (cat #354480) with 8.0 µm PET Membrane in two 24-well Plates. Boyden chamber was rehydrated by adding 400 μL of 1% FCS media to the upper chamber and 700 μL of 1% FCS media to the lower chamber, followed by incubation for 2 h at 37°C in a humidified incubator with 5% CO□. H1975 Par, Osi-R, Bmi1-up and pCMV3 cells were seeded onto the upper chamber, while 700 μL of complete culture medium supplemented with 10% FCS was added to the lower chamber as a chemoattractant. Cells were allowed to migrate for 48 h under standard culture conditions (37°C, 5% CO□). Then, cells were fixed with 1% paraformaldehyde for 10 min at room temperature, washed twice with PBS, and permeabilized with 0.2% Triton X-100 for 10 min. Cells that migrated to the lower surface of the membrane were stained with DAPI (1:1.000) for 10 min. Images were taken with a fluorescence microscope at 5x and 10 x magnification.

### Wound healing assays

H1975-Par and -OsiR cells were seeded in 6 well plates and cultured until a 90-100% confluence was reached. A linear scratch was made across the center of each well using the tip of a sterile 200-μL pipette, ensuring comparable wound widths across replicates. Then, the 10% FBS growth medium was replaced with culture medium supplemented with 2% FBS to minimize cell proliferation during the assay. Images of the wound area were acquired at 0 h, 8 h, 12 h, 14 h, and 16 h. Image processing was performed using the “Wound healing size tool” ImageJ/Fiji® plugin software (Version 1.53c, NIH, Bethesda, MD).

### PTC596/Unesbulin treatment and apoptosis

H1975 Osi-R cells line was treated with Unesbulin (1 μmol/L), or Vehicle (0.5% DMSO) for various time points (24, 48, and 72 hours). Apoptosis was assessed using the Annexin V Apoptosis Detection Kit FITC (Invitrogen, Cat. # 88-88005-74) following the manufacturer’s instructions. Briefly, cells were harvested, filtered through cell strainer caps into FACS tubes (Falcon, Cat. #352235) and resuspended in Annexin V binding buffer 1X. Annexin V master mix (1:20 dilution) was added to each sample and incubated for 10 min in the dark. Cells were washed with binding buffer, centrifuged at 1200 rpm for 5 min at room temperature, and the supernatant was discarded. Propidium iodide (PI) master mix (1:40 dilution) was added. Samples were analyzed withing 4 h by flow cytometry. For each time point the following controls were included:

- A positive control only stained with Annexin V
- A positive control only stained with PI
- A positive control strained with both AnnexinV&PI
- A negative control without AnnexinV and PI staining

FlowJo version 10.0 was used to analyze the apoptosis data.

### Xenograft studies and drug treatments

B-NDG (NOD.CB17-Prkdcscid IL2rgtm1/Bcgen, Envigo) mice were used for xenograft studies. For subcutaneous tumor establishment, 1×10□ H1975-Par, Osi-R, pCMV3-BMI1up or pCMV3 cells were resuspended in 50 μL of Matrigel Basement Membrane Matrix, Phenol Red-free (BD Biosciences, #356237) and injected into both flanks of mice.

Tumor growth was monitored weekly by high-resolution ultrasound imaging beginning on day 10 post-implantation and followed for 3 weeks. To evaluate the growth of the cells *in vivo*, tumor volume (mm^3^) was calculated from ultrasound measurements, and growth kinetics were compared.

To study the effect of Unesbulin *in vivo*, H1975-Osi-R xenografts were randomized to receive Unesbulin (12 mg/kg), or Vehicle (0.5% hydroxypropyl methyl cellulose—HPMC— and 0.2% Tween 80 in distilled water) by oral gavage twice a week, once the tumor became measurable.

For Osimertinib response studies, xenografts with H1975-Osi-R, H1975 Bmi1-up and H1975 pCMV3 cells received biweekly treatment with Osimertinib (10 mg/kg, oral gavage) or vehicle control, once tumors became measurable.

### High Resolution Ultrasound Imaging

Ultrasound (US) imaging was performed using the multimodal imaging platform Vevo LAZR-X system (FUJIFILM VisualSonics Inc., Toronto, Canada) by using high-resolution ultrasound probes. The system is equipped with a sensorized animal bed for physiological monitoring and temperature control. Mice were initially anesthetized in an induction chamber with 2% isoflurane and then moved to the sensorized bed, where anesthesia was maintained via a nose cone with 1% isoflurane. The bed temperature was set around 40 °C throughout the procedure to prevent hypothermia. Depending on the required field of view, two transducers were employed: the X550D probe (frequency range: 25–55 MHz; central band: 40 MHz, axial resolution 40 µm), and the MX250 probe (frequency range: 15–30 MHz; central band: 25 MHz, axial resolution 75 µm).

Tumor morphology was assessed by three-dimensional (3D) imaging of the region of interest using a motorized stepper system (step size: 150 µm), allowing for high-resolution volumetric reconstruction of the tumor area. Coupling between the US probe and tumors was ensured by US clear gel (Cogel, Comedical s.r.l.), ensuring optimal acoustic transmission.

Quantification has been performed using Vevo Lab ultrasound analysis software.

### Statistics

Statistical analyses have been performed using Graphpad Prism 10.4.1 (532). Specific details are provided within each figure legend.

## FUNDING

This work was supported by the AIRC Investigator Grant 2021 (ID 25734), PNNR THE Spoke 1 Award, the PNRRMCNT1-

2023-12377671 Grant, the ELMO Pisa Foundation Grant, the FPS Grant 2024, and private donations from the Gheraldeschi and Pecoraro families to EL.

AC was supported by the European Union - Next Generation EU, Mission 4 Component 2, project “Strengthening BBMRL.it”.

AA acknowledges funding from A-0001263-00-00.

LM acknowledges support from EuroBioImaging-ERIC and the SEELIFE infrastructure funded under the PNRR (Project Code SEELIFE n. IR00023), from Next Generation EU, aligned with the National Recovery and Resilience Plan (PNRR), Mission 4, Component 2, investment number 1.4 and 1.5, within the THE - Tuscany Health Ecosystem (Project Code ECS00000017).”

## Notes

### Competing Interest Statement

The authors have declared no competing interest.

## Bibliography

Benjamini, Y., Drai, D., Elmer, G., Kafkafi, N., and Golani, I. (2001). Controlling the false discovery rate in behavior genetics research. Behav Brain Res 125, 279–284.

Chen, E. Y., Tan, C. M., Kou, Y., Duan, Q., Wang, Z., Meirelles, G. V., Clark, N. R., and Ma’ayan, A. (2013). Enrichr: interactive and collaborative HTML5 gene list enrichment analysis tool. BMC Bioinformatics 14, 128.

Chen, Z., Han, F., Du, Y., Shi, H., and Zhou, W. (2023). Hypoxic microenvironment in cancer: molecular mechanisms and therapeutic interventions. Signal Transduct Target Ther 8, 70.

Fitieh, A., Locke, A. J., Motamedi, M., and Ismail, I. H. (2021). The Role of Polycomb Group Protein BMI1 in DNA Repair and Genomic Stability. Int J Mol Sci 22.

Gupta, P. B., Pastushenko, I., Skibinski, A., Blanpain, C., and Kuperwasser, C. (2019). Phenotypic Plasticity: Driver of Cancer Initiation, Progression, and Therapy Resistance. Cell stem cell 24, 65–78.

Jernigan, F., Branstrom, A., Baird, J. D., Cao, L., Dali, M., Furia, B., Kim, M. J., O’Keefe, K., Kong, R., Laskin, O. L., et al. (2021). Preclinical and Early Clinical Development of PTC596, a Novel Small-Molecule Tubulin-Binding Agent. Mol Cancer Ther 20, 1846–1857.

Kartikasari, A. E. R., Huertas, C. S., Mitchell, A., and Plebanski, M. (2021). Tumor-Induced Inflammatory Cytokines and the Emerging Diagnostic Devices for Cancer Detection and Prognosis. Front Oncol 11, 692142.

Kuleshov, M. V., Jones, M. R., Rouillard, A. D., Fernandez, N. F., Duan, Q., Wang, Z., Koplev, S., Jenkins, S. L., Jagodnik, K. M., Lachmann, A., et al. (2016). Enrichr: a comprehensive gene set enrichment analysis web server 2016 update. Nucleic Acids Res 44, W90–97.

Levantini, E., Maroni, G., Del Re, M., and Tenen, D. G. (2022). EGFR signaling pathway as therapeutic target in human cancers. Semin Cancer Biol.

Maroni, G., Bassal, M. A., Krishnan, I., Fhu, C. W., Savova, V., Zilionis, R., Maymi, V. A., Pandell, N., Csizmadia, E., Zhang, J., et al. (2021). Identification of a targetable KRAS-mutant epithelial population in non-small cell lung cancer. Commun Biol 4, 370.

Maroni, G., Krishnan, I., Alfieri, R., Maymi, V. A., Pandell, N., Csizmadia, E., Zhang, J., Weetall, M., Branstrom, A., Braccini, G., et al. (2024). Tumor Microenvironment Landscapes Supporting EGFR-mutant NSCLC Are Modulated at the Single-cell Interaction Level by Unesbulin Treatment. Cancer Res Commun 4, 919–937.

Meng, X., Wang, Y., Zheng, X., Liu, C., Su, B., Nie, H., Zhao, B., Zhao, X., and Yang, H. (2012). shRNA-mediated knockdown of Bmi-1 inhibit lung adenocarcinoma cell migration and metastasis. Lung Cancer 77, 24–30.

Nishida, Y., Maeda, A., Kim, M. J., Cao, L., Kubota, Y., Ishizawa, J., AlRawi, A., Kato, Y., Iwama, A., Fujisawa, M., et al. (2017). The novel BMI-1 inhibitor PTC596 downregulates MCL-1 and induces p53-independent mitochondrial apoptosis in acute myeloid leukemia progenitor cells. Blood Cancer J 7, e527.

O’Leary, C., Gasper, H., Sahin, K. B., Tang, M., Kulasinghe, A., Adams, M. N., Richard, D. J., and O’Byrne, K. J. (2020). Epidermal Growth Factor Receptor (EGFR)-Mutated Non-Small-Cell Lung Cancer (NSCLC). Pharmaceuticals (Basel) 13.

Oxnard, G. R., Hu, Y., Mileham, K. F., Husain, H., Costa, D. B., Tracy, P., Feeney, N., Sholl, L. M., Dahlberg, S. E., Redig, A. J., et al. (2018). Assessment of Resistance Mechanisms and Clinical Implications in Patients With EGFR T790M-Positive Lung Cancer and Acquired Resistance to Osimertinib. JAMA Oncol 4, 1527–1534.

Ren, H., Du, P., Ge, Z., Jin, Y., Ding, D., Liu, X., and Zou, Q. (2016). TWIST1 and BMI1 in Cancer Metastasis and Chemoresistance. J Cancer 7, 1074–1080.

Ritchie, M. E., Phipson, B., Wu, D., Hu, Y., Law, C. W., Shi, W., and Smyth, G. K. (2015). limma powers differential expression analyses for RNA-sequencing and microarray studies. Nucleic acids research 43, e47.

Sarkar, S., Sahoo, P. K., Mahata, S., Pal, R., Ghosh, D., Mistry, T., Ghosh, S., Bera, T., and Nasare, V. D. (2021). Mitotic checkpoint defects: en route to cancer and drug resistance. Chromosome Res 29, 131–144.

Shapiro, G. I., O’Mara, E., Laskin, O. L., Gao, L., Baird, J. D., Spiegel, R. J., Kaushik, D., Weetall, M., Colacino, J., O’Keefe, K., et al. (2021). Pharmacokinetics and Safety of PTC596, a Novel Tubulin-Binding Agent, in Subjects With Advanced Solid Tumors. Clin Pharmacol Drug Dev 10, 940–949.

Wong, K. Y., Cheung, A. H., Chen, B., Chan, W. N., Yu, J., Lo, K. W., Kang, W., and To, K. F. (2022). Cancer-associated fibroblasts in nonsmall cell lung cancer: From molecular mechanisms to clinical implications. Int J Cancer 151, 1195–1215.

Xie, Z., Bailey, A., Kuleshov, M. V., Clarke, D. J. B., Evangelista, J. E., Jenkins, S. L., Lachmann, A., Wojciechowicz, M. L., Kropiwnicki, E., Jagodnik, K. M., et al. (2021). Gene Set Knowledge Discovery with Enrichr. Curr Protoc 1, e90.

Yang, M. H., Hsu, D. S., Wang, H. W., Wang, H. J., Lan, H. Y., Yang, W. H., Huang, C. H., Kao, S. Y., Tzeng, C. H., Tai, S. K., et al. (2019). Author Correction: Bmi1 is essential in Twist1-induced epithelial-mesenchymal transition. Nature cell biology 21, 533.

Yong, K. J., Basseres, D. S., Welner, R. S., Zhang, W. C., Yang, H., Yan, B., Alberich-Jorda, M., Zhang, J., de Figueiredo-Pontes, L. L., Battelli, C., et al. (2016). Targeted BMI1 inhibition impairs tumor growth in lung adenocarcinomas with low CEBPalpha expression. Sci Transl Med 8, 350ra104.

Zhang, X., Tian, T., Sun, W., Liu, C., and Fang, X. (2017). Bmi-1 overexpression as an efficient prognostic marker in patients with nonsmall cell lung cancer. Medicine (Baltimore) 96, e7346.

